# Vi polysaccharide and conjugated vaccines afford similar early, IgM or IgG-independent control of infection but boosting with conjugated Vi vaccines sustains the efficacy of immune responses

**DOI:** 10.1101/2023.01.03.522561

**Authors:** Siân E. Jossi, Melissa Arcuri, Areej Alshayea, Ruby R. Persaud, Edith Marcial-Juárez, Elena Palmieri, Roberta Di Benedetto, Marisol Pérez-Toledo, Jamie Pillaye, Will M. Channell, Anna E. Schager, Rachel E. Lamerton, Charlotte N. Cook, Margaret Goodall, Takeshi Haneda, Andreas J. Bäumler, Lucy H. Jackson-Jones, Kai-Michael Toellner, Calman A. MacLennan, Ian R. Henderson, Francesca Micoli, Adam F. Cunningham

**Author notes:** These authors contributed equally to this work and share last authorship (Francesca Micoli, Adam F. Cunningham). Francesca Micoli, **Email:**.

## Abstract

Vaccination with Vi capsular polysaccharide (Vi-PS) or protein-Vi typhoid conjugate vaccine (TCV) can protect adults against *Salmonella* Typhi infections. TCVs offer better protection than Vi-PS in infants and may offer better protection in adults. Potential reasons for why TCV may be superior in adults are not fully understood. Here, we immunized wild-type (WT) mice and mice deficient in IgG or IgM with Vi-PS or TCVs (Vi conjugated to tetanus toxoid or CRM_197_) for up to seven months, with and without subsequent challenge with Vi-expressing *Salmonella* Typhimurium. Unexpectedly, IgM or IgG alone were similarly able to reduce bacterial burdens in tissues, and this was observed in response to conjugated or unconjugated Vi vaccines and was independent of antibody being of high affinity. Only in the longer-term after immunization (>5 months) were differences observed in tissue bacterial burdens of mice immunized with Vi-PS or TCV. These differences related to the maintenance of antibody responses at higher levels in mice boosted with TCV, with the rate of fall in IgG titres induced to Vi-PS being greater than for TCV. Therefore, Vi-specific IgM or IgG are independently capable of protecting from infection and any superior protection from vaccination with TCV in adults may relate to responses being able to persist better rather than from differences in the antibody isotypes induced. These findings suggest that enhancing our understanding of how responses to vaccines are maintained may inform on how to maximize protection afforded by conjugate vaccines against encapsulated pathogens such as *S*. Typhi.

## Introduction

*Salmonella enterica* serovar Typhi, *S*. Typhi is the causative agent of typhoid fever which is estimated to cause up to 161,000 deaths/year (1). The high recorded incidence of antibiotic resistance in *S*. Typhi (2), combined with its host restriction to humans, means that improved sanitation and vaccination offer the best routes for control and eradication. There are several licensed vaccines against *S*. Typhi. These are based on the capsular polysaccharide antigen Vi (3), or the live attenuated strain Ty21a (4, 5). Purified, unconjugated Vi polysaccharide (Vi-PS) and Ty21a offer similar levels of protection in the first two years post-vaccination, but neither are licensed for use in infants, a population particularly affected by typhoid (6-9), nor is the protection they offer sustained for more than a few years (10, 11).

In recent years, there have been major efforts to develop glycoconjugate vaccines against *S*. Typhi, which can induce effective immune responses in both infants and adults. The first conjugate tested, using enterotoxin A from *Pseudomonas aeruginosa* as a carrier protein, demonstrated excellent efficacy in the field (12). This is likely to be because conjugation of Vi-PS to a carrier protein can effectively convert T-independent responses to the polysaccharide antigen into T-dependent responses. The presence of a conjugated protein moiety enables the use of the vaccine in all age groups, and the ability to boost responses through the induction of memory. Studies have shown that boosting with T-independent antigens like Vi vaccine does not augment antibody responses and may even induce hypo-responsiveness (13-15). Newer generations of Typhoid Conjugate Vaccines (TCVs) have been developed using flagellin or classical carrier proteins such as tetanus toxoid (TT) (16, 17) or diphtheria toxoid (18, 19) and its modified form CRM_197_ (20). In most cases, TCVs have shown promise in animal and human challenge studies as well as field trials (21-23), leading to the recent pre-qualification for use by the World Health Organization’s Strategic Advisory Group of Experts (SAGE) of two typhoid TCVs, Typbar and TyphiBev (24, 25).

In a recent study, we investigated the response in mice to TCVs of different compositions and obtained through different conjugation strategies (26). These studies showed that conjugation of Vi to different carrier proteins had little effect on immunogenicity. What was not assessed in these studies was the relationship between immunogenicity, antibody isotype, protection, and the longevity of immunity. This is important to understand, as why the reasons why TCVs are effective in infants are clear, yet the mechanistic basis of why TCVs may provide enhanced protection compared to Vi-PS in adults is incompletely understood. One limitation in achieving this is the human restriction of *S*. Typhi and combined with this being a category III pathogen makes working with this organism restrictive. In part, such difficulties can be overcome through engineering *Salmonella* Typhimurium to express Vi antigen (27-29). We and others have shown that such engineered pathogens can colonize mice and we have previously shown that antibodies induced to Vi vaccines are sufficient to reduce bacterial burdens in mice (28). In this study, we have addressed the response to Vi-PS and TCV in wild-type (WT) mice or mice deficient in specific elements of the antibody response. Additionally, in some experiments mice have been challenged with Vi-expressing *S*. Typhimurium to assess the capacity of antibodies to different vaccine types to reduce bacterial burdens in tissues. The results from these studies indicate there is significant redundancy in the nature of protective antibodies that target the Vi capsule but identify that maintaining the longevity of responses plays a key role in protection.

## Materials and Methods

### Antigens

Vi-CRM_197_ and Vi-TT conjugates were synthesized as previously described (25). For generation of Vi-biotin, fragmented Vi (fVi) with an average MW of 43KDa was used (25). fVi was solubilized in 400 mM MES pH 6 at a concentration of 50 mg/mL. NHS was added to a final concentration of 0.33 M. EDAC was added to a molar ratio of 1:5 with respect to fVi. The reaction was mixed at room temperature for 1 hour. The biotin dihydrazide linker was added with 1:1 w/w ratio with respect to fVi, at a fVi concentration of 40.4 mg/mL. The linker was solubilized in DMSO to a final DMSO concentration of 18.9%. The reaction was mixed for 2 hours at room temperature (RT) and then 4M NaCl was added to the derivatized fVi for purification by PD-10 column (GE Healthcare) against water. Derivatization was confirmed by 1H-NMR analysis showing 13.5% of Vi repeating units were linked to biotin.

### Bacterial strains and growth conditions

Vi+ *S*. Typhimurium TH177 (29) was grown on LB agar plates, or in LB broth supplemented with 30 µg/mL chloramphenicol at 37 °C. For mouse infection, overnight cultures were re-grown to OD_600_ 0.8 in LB without antibiotics at 37 °C, 180 rpm, and washed 3x with PBS. Bacterial concentration was determined by serial dilution on LB agar plates.

### Animal studies

C57Bl/6 mice were sourced from Charles River Laboratories. AID^-/-^ (30), HMT (IgG1^-/-^) (31), and IgM^s-/-^ mice (32) on a C57Bl/6 background were bred in the University of Birmingham Biomedical Service Unit. Both male and female mice were used at age 6-12 weeks. All procedures were performed with ethical approval from the University of Birmingham and UK Home Office approval in compliance with the Animals (Scientific Procedures) Act 1986.

Immunization was performed by intraperitoneal (*i.p*.) injection with Vi-PS or Vi-conjugate diluted in 200 µL PBS with doses indicated in the text referring to the amount of Vi antigen given. Concentration was measured as Vi content, via High Performance Anion Exchange Chromatography with Pulsed Amperometric Detection (HPAEC-PAD) (33). Where stated, a further booster dose was administered via the same route. Bacterial challenge was performed by *i.p*. administration of 1×10^5^ CFU S. Typhimurium TH177 in 200 µL PBS.

### Serum Enzyme-Linked Immunoassays (ELISA)

ELISA plates were coated with 5 µg/mL purified Vi at 4 °C overnight. Wash steps were performed with PBS 0.05 % tween-20 (v/v; PBS-T). PBS 1 % BSA (w/v) was used as a blocking buffer. Incubation steps were performed for 1 hour at 37 °C. Mouse serum was obtained by centrifugation of clotted whole blood at 6000 rpm, then serially diluted 1:3 from a starting dilution of 1:50 or 1:100 into PBS-T 1 % BSA. Anti-Vi antibody was detected by AP-conjugated goat anti-mouse secondary antibodies (Southern Biotech) at 1:4000 for IgM, 1:2000 for IgG2b, and 1:1000 for all other IgG. Secondary antibody was detected with Sigma FAST p-NPP tablets (Sigma Aldrich), and optical density (OD) read at 405 nm using an SpectraMax ABS Plus plate reader (Molecular Devices). Relative reciprocal titres were calculated by measuring the dilution at which serum reached a defined OD_450_.

### Enzyme-Linked ImmunoSpot (ELISPOT) assay

To obtain bone marrow cells, the femur and tibia of one leg per mouse was cleaned and marrow flushed out using a 25 µM gauge needle and 3 mL R-10 media (RPMI 1640 media supplemented with 10 % heat-inactivated fetal bovine serum and 5 % penicillin-streptomycin). Splenocytes were obtained by disruption of the spleen tissue through a 70 µM cell strainer into R-10. For both tissue suspensions, erythrocytes were lysed with 500 µL ammonium-chloride-potassium (ACK) lysis buffer. Cells were centrifuged at 1400 xg and washed in R-10 before and after lysis.

Multiscreen filter plates (Merck Millipore) were ethanol treated for 1 minute and washed with PBS before coating with 5 µg/mL purified Vi at 4 °C overnight. Plates were washed and blocked with R-10 media for 2-3 hours at 37 °C. Wells were washed and seeded with 5×10^5^ cells/well and incubated for 6 hours at 37 °C, 5 % CO_2_. Cells were washed away 3x with PBS 0.05 % tween and 1x with PBS. The remaining bound anti-Vi antibody was detected by AP-conjugated goat anti-mouse secondary antibodies (Southern Biotech) at 1:1000 in PBS at 4 °C overnight. Plates were washed as before, and spots developed with Sigma FAST BCIP tablets (Sigma Aldrich). Once the plates were dried, spots were detected using an AID iSPOT Spectrum plate reader (Advanced Imaging Devices GMBH) and counting adjusted by eye.

### Immunohistochemical staining (IHC)

Spleens were frozen in liquid nitrogen and sectioned at 6 µM for transfer to glass slides. Slides were air dried and fixed in acetone for 15 minutes. All antibodies were prepared in 75 µL Tris-HCl pH 7.6 and incubated in a humidity chamber for 1 hour at room temperature. Peroxidase labelled rat anti-mouse IgD (Southern Biotech), IgM (AbD Serotec) and IgG (BioRad) antibodies were used at dilutions 1:1000, 1:500 and 1:300 respectively. Biotinylated PNA (Vector Laboratories) was used at 1:200, and Biotinylated Vi used at a 1:150 dilution. Signal was developed with 3,3-diaminobenzidine tablets (Sigma Aldritch). Biotinylated reagents were detected with a streptavidin-biotinylated AP complex (Vector Laboratories) and developed with Naphthol AS-MX phosphate, levamisole, and Fast Blue Salt.

### Bacterial burden determination

To measure bacterial load of mouse spleens and livers, the tissues were passed through a 70 µM cell strainer into 1mL PBS. The cell suspension was serially diluted 1:10 and spread as a lawn on LB agar plates. Plates were incubated at 37 °C for ∼16 hours before colonies were counted. Colony counts were converted to colony forming units (CFU) per gram of tissue used.

### Serum Bactericidal Assay (SBA)

To determine serum bactericidal activity, sera were heat inactivated for 30 minutes at 56 °C for complement inactivation. Human serum adsorbed against *S*. Typhimurium TH177 was added at a 1:1 ratio to heat-inactivated mouse serum samples as a complement source (34, 35). Complete human serum (CHS) was used as a positive control for bactericidal activity. Then 2×10^7^ CFU *S*. Typhimurium TH177 was added to the serum samples and incubated at 37 °C with gentle rotation. Ten µL was removed at after 45, 90 and 180 mins and serially diluted into PBS before plating onto LB agar plates in triplicate. Plates were incubated overnight at 37 °C and colonies counted.

### IgM depletion from mouse sera

To remove IgM from mouse sera, either serum from one naïve mouse, or pooled sera from mice immunized and boosted three times with Vi-TT were used. Rat polyclonal anti-mouse IgM antibody was conjugated onto cyanobromide activated Sepharose beads (Sigma Aldritch) as per the manufacturer’s instructions. 200 µL of serum was incubated with 200 µL of antibody-conjugated beads for 30 minutes with gentle shaking. Samples were centrifuged at 2000 rpm and supernatants transferred to a clean tube. Beads were washed twice with sterile PBS and once with 0.8 M pH 2.0 citric acid to eluate bound IgM. Four rounds of depletion were performed until IgM was undetectable by ELISA. Total IgG1 was also detected to quantify loss of antibody due to dilution in the column alone. ELISAs were conducted as described above, using 1 µg/mL of rat anti-mouse IgM (AbD Biotec) or rabbit anti-mouse IgG1 (Dako) capture antibody, or purified Vi as a target.

### Opsonization of live bacteria

For opsonization experiments, bacteria were prepared as described above to the desired infection dose in 900 µL of sterile PBS. Sera was added to the bacterial dose to a final dilution of 1:100. Samples were then incubated at room temperature with gentle agitation for 30 minutes before use.

### Statistical analysis

Statistical analysis was performed using GraphPad Prism 8.0. Where samples passed the D’Agostino & Pearson test for normality, an unpaired t-test was used to compare two groups and an ordinary one-way or two-way ANOVA for groups >2. Where n was too small or a non-normal distribution was observed, a Mann-Whitney U test or a Kruskall-Wallace ANOVA with Dunn’s post-hoc multiple comparisons test were used, respectively. Significance was accepted as p≤0.05.

## Results

### Immunization with TCVs induces rapid, higher Vi-specific IgG responses than Vi-PS

To compare the antibody responses induced by conjugated and non-conjugated Vi vaccines, WT C57BL/6 mice were immunized intraperitoneally with 2 µg of Vi-PS, Vi-TT or Vi-CRM_197_. After 7 days, all vaccines induced Vi-specific serum IgM and IgG antibody, but both TCVs induced higher titres of each (Figure 1A). Frequencies of Vi-specific antibody secreting cells (ASCs) amongst splenocytes were analyzed by ELISPOT, and at 7 days post-immunization TCVs induced greater median IgG+ Vi-specific ASC responses than Vi-PS (Figure 1B). Moreover, immunohistology identified that Vi-specific plasmablasts/cells (PB/C) localized to classical extrafollicular foci in the red pulp of the spleen after immunization with Vi-PS and TCVs (Figure 1C). Both IgM+ and IgG+ Vi-specific ASCs were detected (Supplementary Figure 1A). Germinal centers (GCs) above background levels were only detectable in mice immunized with Vi-conjugate (Figure 1C). Analysis of the immunoglobulin switching pattern induced by the vaccines at day 7 showed that all Vi vaccines induced predominantly IgG1 and IgG3 (Figure 1D). At later time-points, in addition to IgG1 and IgG3, IgG2b was also detected consistently. Some IgA was detected after immunization with TCV but not Vi-PS, but these responses were only modestly above background and did not persist over time in most mice (Supplementary Figure 1B). Thus, after immunization with Vi vaccines, a rapid IgM and IgG extrafollicular response is induced to all Vi vaccines, but a notable GC response is only seen against TCVs.

**Figure 1.**
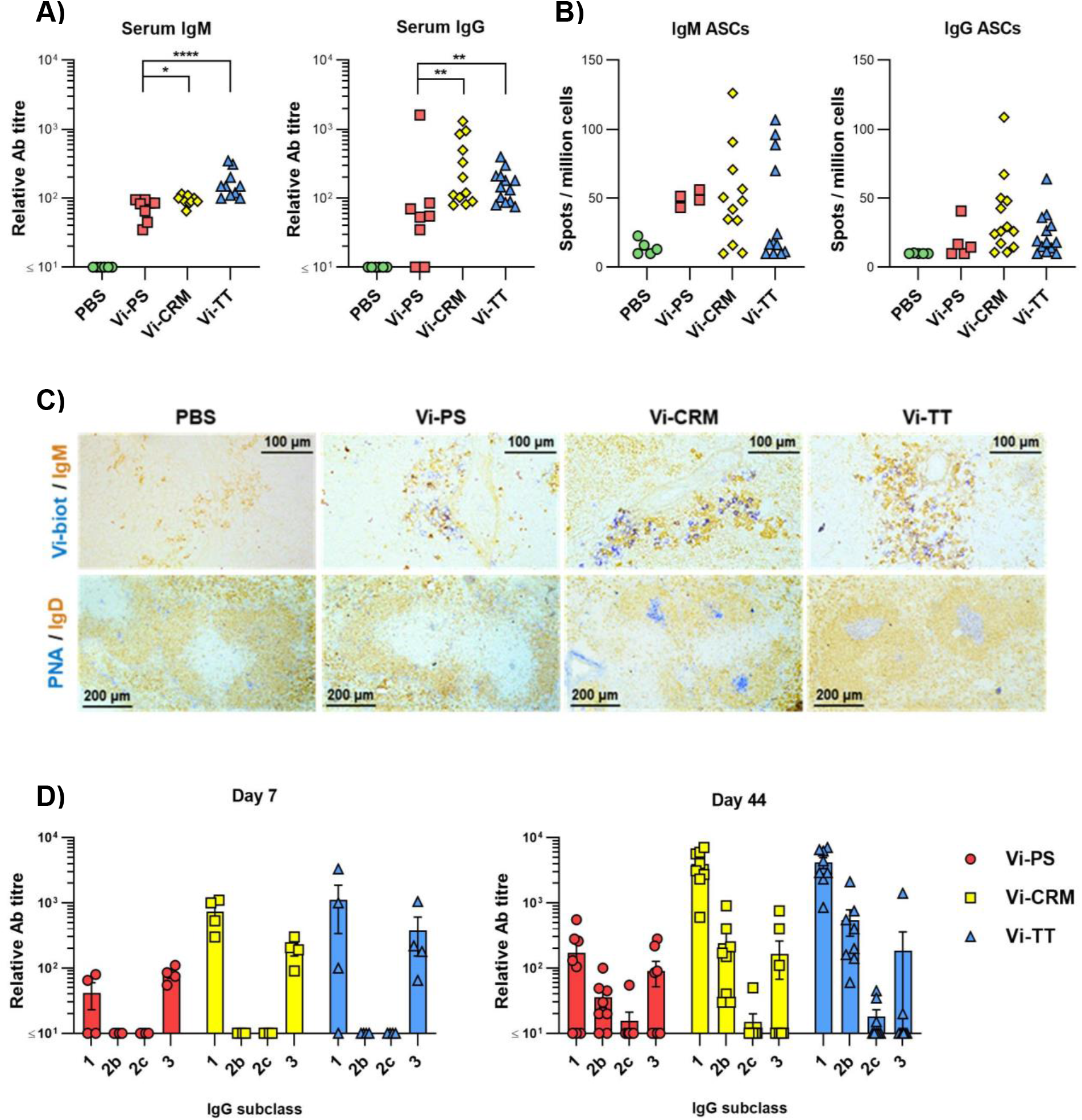
TCV induce rapid IgM and IgG responses and GC. **(A)** Sera from C57Bl/6 mice 7 days after *i.p* injection of 2 µg Vi-PS, Vi-CRM_197_, Vi-TT or PBS were assessed by ELISA for anti-Vi IgM and IgG. Data pooled from 3 experiments with n = 2 or 4 mice/group. **(B)** Splenic antibody secreting cells (ASCs) producing anti-Vi IgM or IgG were enumerated by ELISPOT, with representative well images below each group. **(C)** Representative immunohistological images of spleen tissue stained for IgM (brown) and Vi (blue), or IgD (brown) and PNA (blue). 10x magnification. **(D)** ELISA was used to quantify anti-Vi IgG1, IgG2b and IgG3 in sera from mice 7 days post-immunization, or sera from C57Bl/6 mice immunized *i.p* with 2 µg Vi-PS, Vi-CRM_197_, Vi-TT or PBS on day 0 and 35, then challenged with 1×10^5^ CFU Vi+ *S*. Typhimurium TH177 from day 41-44. Each timepoint is composed of 2-3 experiments with n = 2 or 4 mice/group. Bars represent mean relative antibody titre of each isotype, with SEM. * = p≤0.05, ** = p≤0.01 and **** = p≤0.001 by Mann-Whitney U test between individual groups (two-tailed).

### Vi-PS and TCVs offer similar early protection against infection

To determine if immunization with TCVs resulted in reduced bacterial burdens in tissues after infection compared to Vi-PS vaccines, mice were immunized with Vi-PS or either TCV, then challenged 14 days later with Vi-expressing *S*. Typhimurium strain TH177 (Figure 2A). All vaccines induced similar reductions in bacterial burdens in the spleen compared to non-immunized mice, with no significant difference in the protection afforded by the different vaccines. Bacterial numbers were also assessed in mice that were immunized once with Vi-PS or Vi-TT (4 µg at day 0) or twice with Vi-PS, Vi-CRM, or Vi-TT (2 µg each at days 0 and 35). In these mice, anti-Vi titres were increased after a second dose of Vi-conjugate, but no such booster effect was observed for Vi-PS (Supplementary Figure 2). Similar bacterial burdens were observed in all the immunized and challenged groups (Figure 2B). Therefore, both Vi-PS and TCVs induce similar protection in the first few weeks after immunization, independent of whether mice are immunized once or are boosted with Vi-PS or TCV.

**Figure 2.**
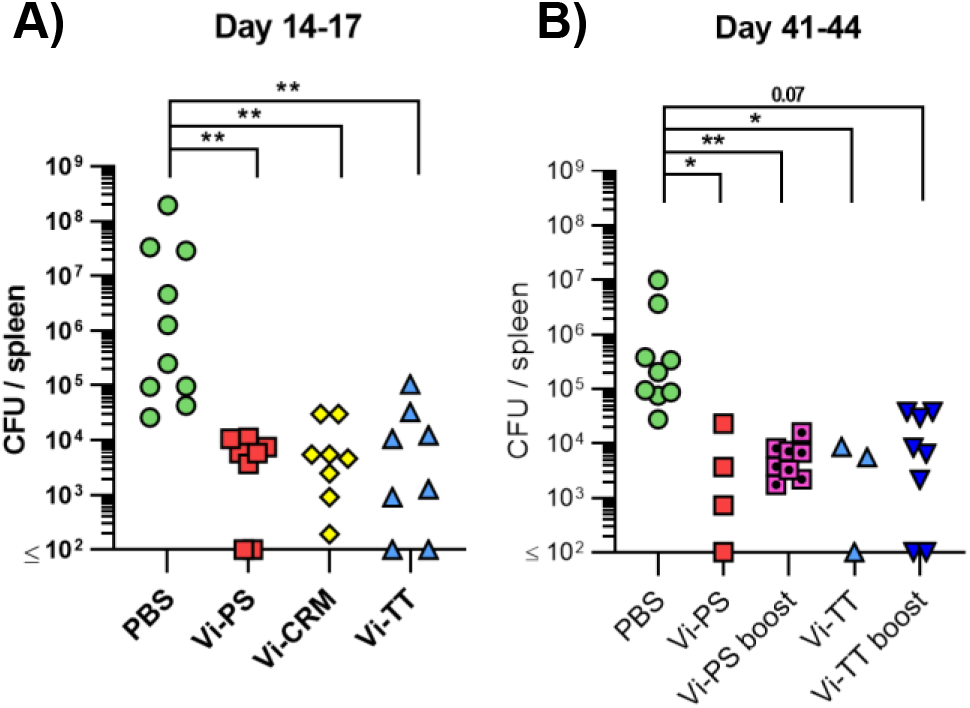
Vi-polysaccharide and Vi-conjugate vaccines provide equal protection against infection with Vi+ *S*. Typhimurium. **(A)** Colony forming units (CFU) in the spleen of C57Bl/6 mice immunized *i.p* with 2 µg Vi-PS, Vi-CRM_197_, Vi-TT or PBS mice, then infected for 72 hours with 1×10^5^ CFU Vi+ *S*. Typhimurium TH177 14 days later. Data pooled from 2 experiments with n = 4 mice/group. **(B)** Spleen CFU of immunized *i.p* with either 4 µg of Vi-PS or Vi-TT at day 0, or 2 µg each vaccine at both day 0 and 35 (boost groups), then infected with 1×10^5^ CFU TH177 day 41, for 72 hours. Data pooled from 2-3 experiments with n = 2 or 4 mice/group. * = p≤0.05, ** = p≤0.01, *** = p≤0.005 and **** = p≤0.001 by Kruskall Wallace one-way ANOVA with Dunn’s multiple comparisons between groups (two-tailed).

### Anti-Vi IgM induced by Vi vaccines is sufficient to impair infection

In previous work, an important role for IgG1 against STm after immunization with the bacterial porin OmpD was identified (36). To test whether IgG1 was similarly important for the protection afforded by immunization with Vi, mice lacking IgG1 (HMT mice) were immunized once with Vi-PS or Vi-TT and challenged with STm TH177 14 days later. As both Vi-TT and Vi-CRM_197_ induced similar responses, Vi-TT was selected as a representative TCV in these experiments. Both Vi vaccines induced similar levels of anti-Vi IgM and total IgG (Supplementary Figure 3A) and upon challenge both immunized groups had similar reductions in the bacterial numbers recovered (Figure 3A).

**Figure 3.**
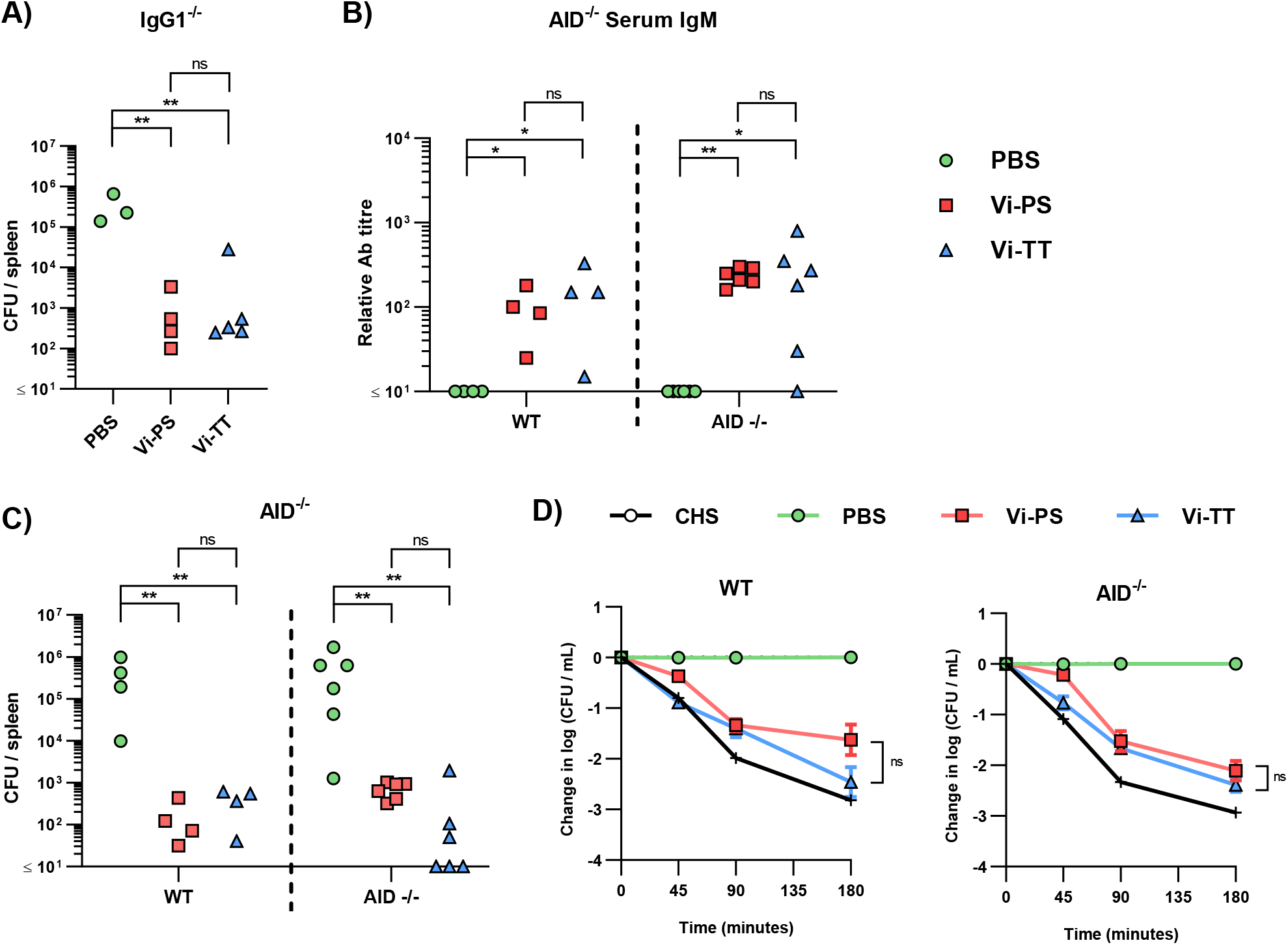
Anti-Vi IgM induced by Vi vaccines is sufficient to impair infection. **(A)** Colony forming units (CFU) in the spleen of IgG1^-/-^ mice immunized *i.p* with 2 µg Vi-PS, Vi-TT or PBS, then infected for 24 hours with 1×10^5^ CFU Vi+ *S*. Typhimurium TH177 14 days later. Representative of 2 experiments. **(B)** Serum anti-Vi IgM titres of WT and AID^-/-^ mice after the same challenge schedule were assessed by ELISA after absorption against BSA. **(C)** Spleen CFU counts of both C57B/6 (WT) and AID^-/-^ mice. **(D)** Serum bactericidal activity of WT and AID^-/-^ sera was measured against TH177, represented as mean change from the starting LogCFU/mL bacterial dose, normalised to the negative control values at each timepoint. Complete human serum (CHS) was used as a positive control. Error bars represent SEM. Representative of 2 experiments with n = 2-4 mice/group. * = p<0.05, ** = p<0.01, and ns = non-significant by Mann-Whitney U test between individual groups (two-tailed).

The lack of a major role for IgG1 in protection suggested that after immunization with TCV other IgG subclasses may be able to compensate for the loss of IgG1. To test this, WT and mice deficient in AID, which cannot switch to any IgG class (37), were immunized for 14 days with Vi-PS or Vi-TT and then challenged with STm TH177. As expected, no Vi-specific IgG was detected in AID^-/-^ mice, irrespective of which vaccine was given, but Vi-specific IgM titres were similar between WT and AID^-/-^ mice (Figure 3B; Supplementary Figure 3B). Immunized WT or AID^-/-^ mice had similar bacterial burdens after challenge, independent of which vaccine type they received (Figure 3C). The sera obtained after immunization were tested in a complement-dependent serum bactericidal assay (34). Sera from WT or AID^-/-^ mice that received either vaccine could promote killing (Figure 3D). Thus, for both Vi-PS and TCVs when IgM is present at comparable levels between the groups then IgM alone is sufficient to reduce bacterial numbers *in vitro* and *in vivo*.

### Anti-Vi IgG induced by Vi vaccines is sufficient to impair infection

The above experiments suggest that IgM to Vi is sufficient or essential to control bacterial burdens after immunization with different Vi vaccines. To determine this, mice that express surface IgM, but that do not secrete IgM (IgM^s-/-^ mice), were immunized with Vi-PS or Vi-TT and challenged 14 days later with STm TH177. All immunized mice had reduced bacterial burdens and these bacterial burdens were similar between WT and IgM^s-/-^ mice (Figure 4A). No IgM was detectable in the serum of IgM^s-/-^ mice (Supplementary Figure 3C), but anti-Vi IgG was detected in all immunized mice, albeit at higher titres in mice immunized with Vi-TT (Figure 4B). Testing the sera from the different groups in the SBA showed that sera derived from immunized WT and IgM^s-/-^ mice were able to kill bacteria (Figure 4C), with bacterial numbers least reduced when sera from Vi-PS IgM^s-/-^ mice were tested. To confirm these results, we used beads coated with rat anti-mouse IgM monoclonal antibodies to deplete IgM from hyperimmune anti-Vi sera generated by immunizing three times with Vi-TT (Supplementary Figure 4). IgM-depleted serum was as capable of killing STm TH177 in the SBA as the same serum prior to depletion (Figure 4D). Moreover, mice infected with STm TH177 bacteria opsonized with IgM-depleted or non-depleted sera had similarly reduced bacterial burdens (Figure 4E). Thus, anti-Vi IgG is capable of driving protection from infection, independent of whether it is induced by Vi-PS or TCV.

**Figure 4.**
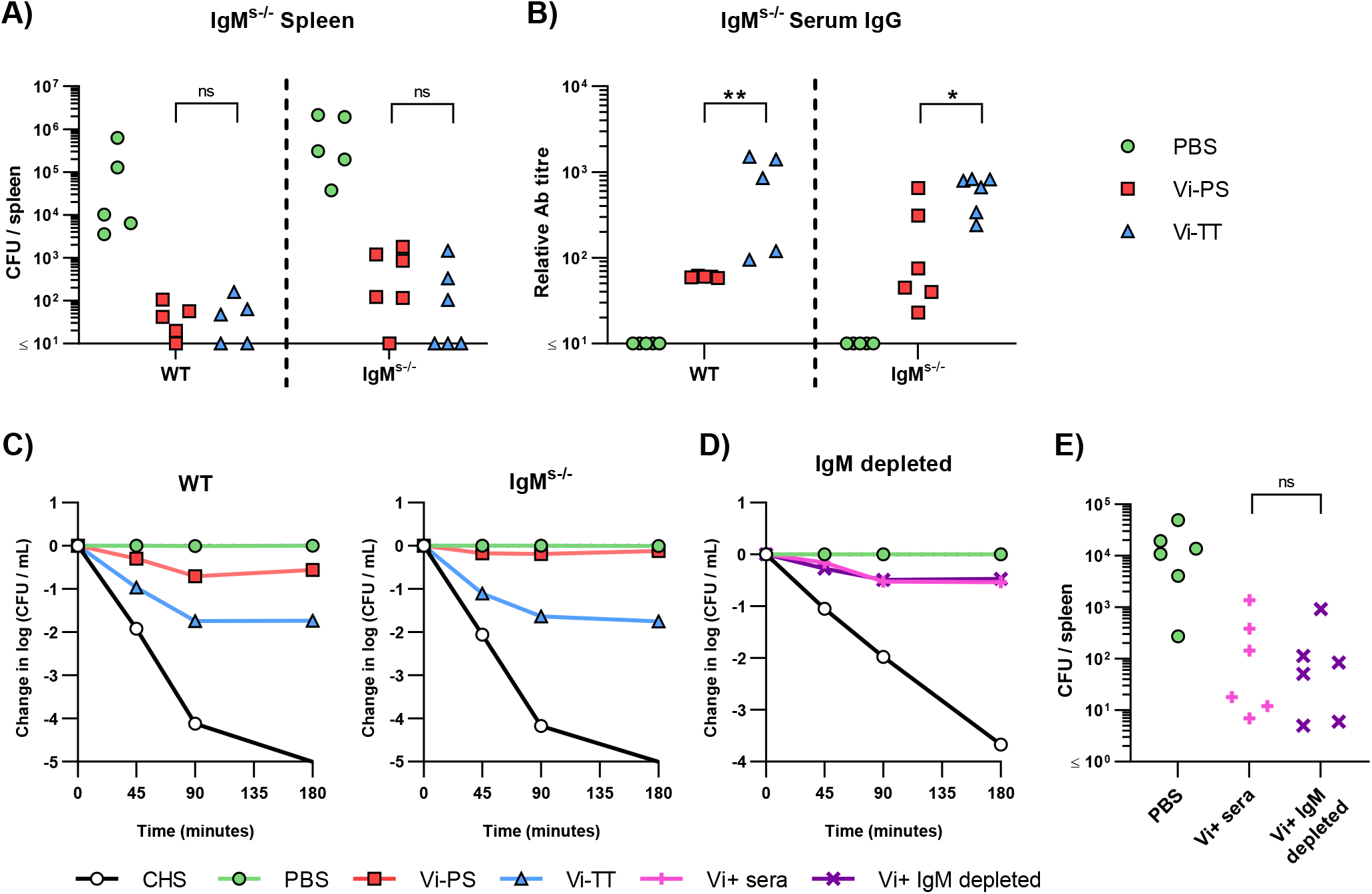
Anti-Vi IgG induced by Vi vaccines is sufficient to impair infection. **(A)** Colony forming units (CFU) in the spleen of C57Bl/6 (WT) and IgM^s-/-^ mice immunized *i.p* with 2 µg Vi-PS, Vi-TT or PBS, then infected for 24 hours with 1×10^5^ CFU Vi+ *S*. Typhimurium TH177 14 days later. Data pooled from 2 experiments with n = 2 or 4 mice/group. **(B)** Serum anti-Vi IgG titres of WT and IgM^s-/-^ mice were assessed by ELISA. **(C)** Serum bactericidal activity of WT and IgM^s-/-^ sera was measured against TH177, represented as mean change from the starting LogCFU/mL bacterial dose, normalised to the negative control values at each timepoint. Error bars represent SEM. **(D)** Pooled WT hyperimmune anti-Vi sera were depleted of IgM using anti-mouse IgM coated Sepharose beads, and serum bactericidal activity against TH177 measured by SBA. Representative of individual serum samples, showing change from the starting LogCFU/mL bacterial dose, normalised to the negative control values at each timepoint. Complete human serum (CHS) was used as a positive control. **(E)** Spleen CFU counts of WT mice infected for 24 hours with 1×10^5^ CFU TH177 opsonised with IgM depleted serum. Data pooled from 2 experiments with n= 2 or 4 mice/group. * = p<0.05, ** = p<0.01 and ns = non-significant by Mann-Whitney U test between individual groups (two-tailed).

### Antibody responses to TCVs persist at higher levels than those to Vi-PS

Since the presence of anti-Vi IgM or IgG induced by either vaccine was sufficient to protect from infection, we hypothesized that in adults TCVs are more effective than Vi-PS vaccines (38-41) because boosting induces longer-lasting antibody responses to the same total dose of Vi. To test this, WT mice were immunized with either 4 µg of Vi-PS once, or 2 µg of TCV twice, 35 days apart. Serology at 6 months after vaccination showed that nearly all immunized mice had serum IgG at this time and the IgG titres were significantly higher in mice immunized twice with TCV than immunized with Vi-PS (Figure 5A). The isotype switching pattern observed at this late time-point reflected that seen earlier in the response (Figure 5B). In addition, nearly all mice boosted with a TCV also had detectable levels of IgM (Figure 5A), which was not the case after immunization with Vi-PS. Anti-Vi IgM was not a consistent feature of the response to TCV in mice immunized once with 2 µg of TCV 6 months previously, despite IgG responses being maintained in most mice receiving this lower dose of vaccine (Supplementary Figure 5A). Thus, at 6 months post-immunization, serum Vi-specific antibody levels are higher in mice receiving two doses of 2 µg of TCV, compared to immunization with 4 µg of Vi-PS.

**Figure 5.**
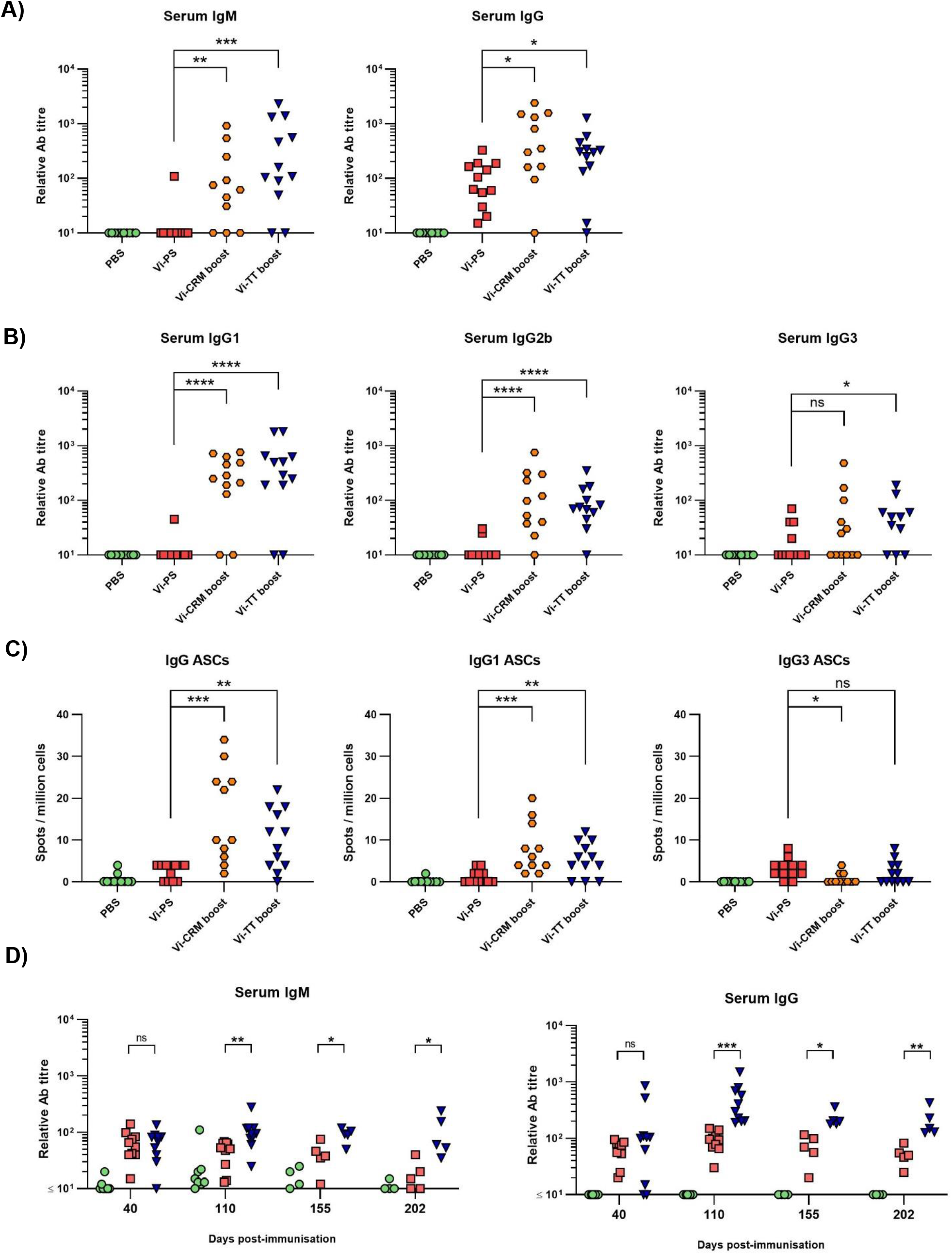
Antibody responses to Vi-PS and TCVs over time. C57Bl/6 mice were immunized *i.p* with PBS, 4 µg Vi-PS on day 0, or 2 µg of Vi-CRM_197_ or Vi-TT on both day 0 and 35. Serum was collected day 185 (6 months post-primary immunization). Sera were assessed by ELISA for **(A)** anti-Vi IgM and IgG and **(B)** isotypes IgG1, IgG2b or IgG3. **(C)** Bone marrow anti-Vi IgG, IgG1 and IgG3 antibody secreting cells (ASCs) at the 6-month timepoint were enumerated by ELISPOT. **(D)** C57Bl/6 mice were immunized *i.p* with 2 µg Vi-PS or PBS on day 0, or 2 µg Vi-TT at both day 0 and 35. Serum was collected day at day 40, 110, 155 and 202 and anti-Vi IgM and IgG determined by ELISA. Data pooled from 2 experiments with n = 5-6 mice/group. * = p≤0.05, ** = p≤0.01, *** = p≤0.005, **** = p≤0.001 and ns = non-significant by Mann-Whitney U test between individual groups (two-tailed).

A proxy indication for the persistence of vaccine-induced responses are the frequencies of antigen-specific antibody-secreting cell (ASC) in niches such as the bone marrow (BM). At 6 months, Vi-specific IgG ASCs were detectable in >95% of mice that received two doses of 2 µg of TCV and only in 50% of mice immunized with 4 µg of Vi-PS (Figure 5C). Moreover, the frequencies of ASC in mice immunized twice with Vi-TT were higher than Vi-PS-immunized mice. Even in mice immunized with one dose of 2 µg of TCV >85% of mice had detectable IgG ASC in the BM (Supplementary Figure 5B). IgG1-secreting ASC and IgG3 were most readily detected after immunization (Figure 5C).

We then examined how IgM and IgG responses to the different vaccines persist over time. To do this, sera were collected at 40, 100, 155 and 202 days after immunization with either Vi-PS (4 µg) or two doses of Vi-TT (2 µg, 35 days apart) and assessed for Vi-specific IgM and IgG (Figure 5D). At day 40, the differences in titres between the immunized groups were only modest, <2-fold different. By day 100 onwards they were consistently greater than 3-4-fold higher in mice receiving Vi-TT than Vi-PS. We compared the differences in IgM and IgG titres between days 110 and day 202 as a measure of the rate of decay of the circulating antibody response. At these times, it is likely that the generation of new plasma cells induced by the vaccination is complete (42-44). In this ∼100-day timeframe, mice immunized with Vi-PS or Vi-TT saw a similar decline in median anti-Vi IgM titres of approximately 47 and 58% respectively (Supplementary Figure 6). However, while anti-Vi IgG titres in Vi-PS immunized mice fell significantly during this time (median 71% fall), IgG titres in the Vi-TT group did not (median 36% fall). These results suggest that boosting can enhance the maintenance of anti-Vi IgG antibody responses through inducing higher IgG responses overall and by resulting in a slower decline in these responses thereafter.

**Figure 6.**
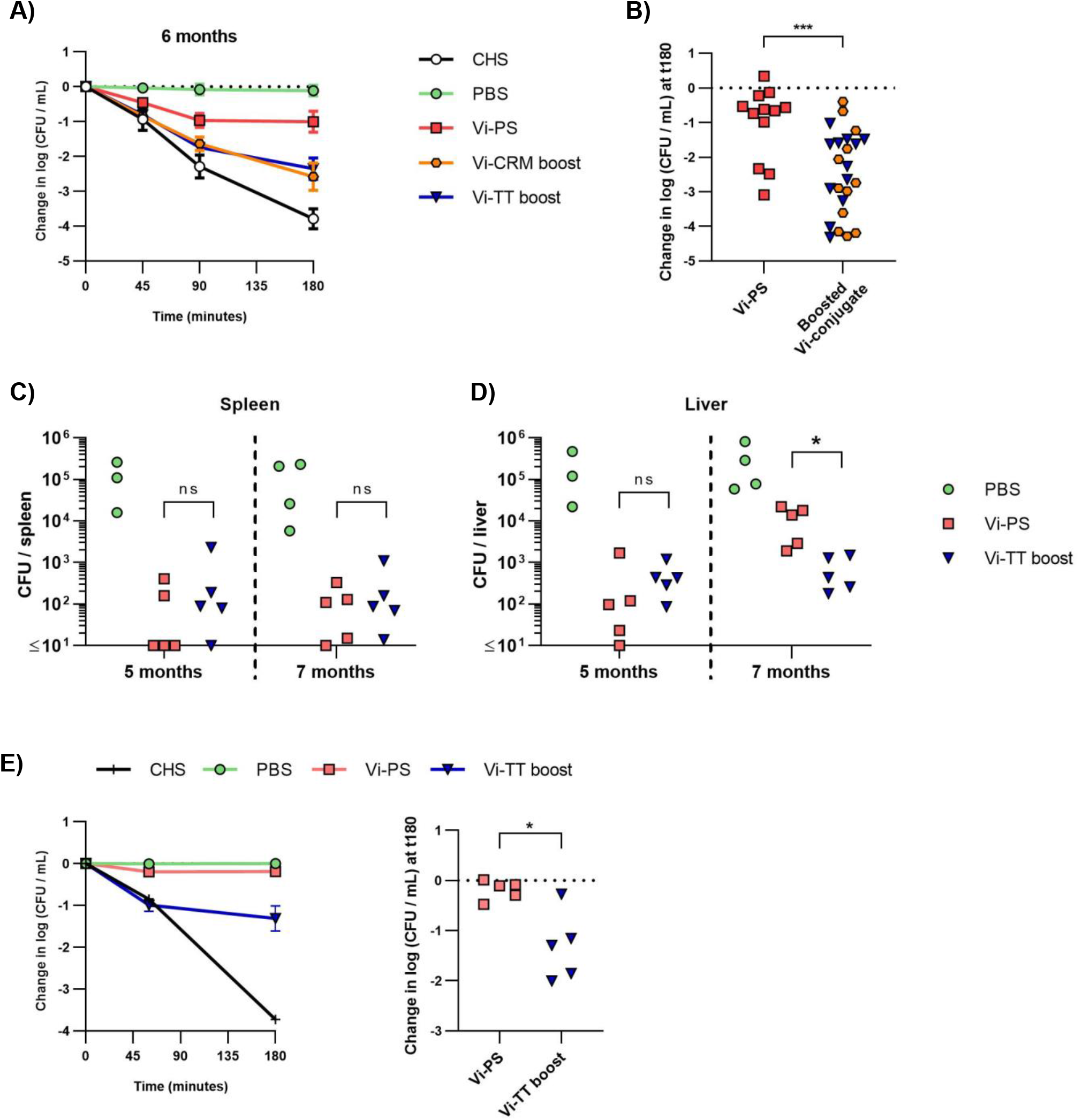
Bactericidal activity of mice immunized with Vi vaccines over time and the effect on protection. C57Bl/6 mice were immunized *i.p* with PBS, 4 µg Vi-PS, on day 0, or 2 µg Vi-CRM_197_ or Vi-TT at both day 0 and 35. **(A)** Serum bactericidal activity of sera collected day 185 (6 months post-primary immunization) was measured against TH177, represented as mean change from the starting LogCFU/mL bacterial dose over time, normalised to the negative control values at each timepoint. Complete human serum (CHS) was used as a positive control. Error bars represent SEM. **(B)** Serum bactericidal activity from the 180-minute timepoint represented as mean change from the starting LogCFU/mL between mice immunized once at day 0 with Vi-PS. or twice at day 0 and 35 with Vi-CRM_197_ or Vi-TT. **(C)** Spleen or **(D)** Liver colony forming units (CFU) of C57Bl/6 (WT) mice immunized *i.p* with 2 µg Vi-PS or PBS on day 0, or 2 µg Vi-TT at both day 0 and 35 then infected for 24 hours with 1×10^5^ CFU Vi+ *S*. Typhimurium TH177 at day 154 or 201 (5 or 7 months post-primary immunization). Representative of 2 experiments with n = 4 or 5 mice/group. For b-d each point represents data from one mouse. **(E)** Serum bactericidal activity of sera taken day 199 (prior to day 202 infection), represented as change from the starting LogCFU/mL bacterial dose normalised to the negative control values at each timepoint, or at the 180 minuet timepoint alone. Complete human serum (CHS) was used as a positive control. Groups contain 4-5 mice/group and in the right hand graph each point represents one serum. * = p≤0.05, ** = p≤0.01, *** = p≤0.005, **** = p≤0.001 and ns = non-significant by Mann-Whitney U test between individual groups (two-tailed).

### Functional antibody responses are reduced after long-term immunization with Vi-PS

Finally, we examined the capacity of long-term immunized mice to control infection and of sera from these mice to kill bacteria in the SBA. Sera obtained from the 6-month immunized mice (Figure 5) were evaluated for their activity in SBA (Figure 6A-B). Sera from all immunized mice had some killing activity compared to non-vaccinated mice, even when Vi-specific antibodies were present in low titres. The bactericidal activity was significantly lower in sera from mice immunized with Vi-PS than in sera from mice given two doses of TCV (Figure 6B). To test if the differences in SBA activity reflected the capacity of these mice to control infection *in vivo*, mice were immunized with a single 4 µg dose of Vi-PS, or two 2 µg doses of Vi-TT (35 days apart) and challenged with STm TH177 at either 5 or 7 months post-primary immunization. At both 5- and 7-months post-immunization, the bacterial burden in the spleens of Vi-PS and Vi-TT immunized mice were significantly lower than for non-immunized mice (Figure 6C). Moreover, at 5 and 7 months after challenge, mice immunized with Vi-TT had similar splenic and liver bacterial burdens respectively, suggesting a similar level of protection is maintained between these time-points (Figure 6D). Mice immunized with Vi-PS had similar bacterial burdens as Vi-TT-immunized mice except that bacterial burdens were higher in the liver at 7 months (Figure 6D), suggesting vaccine efficacy in Vi-PS-immunized mice is only modestly reduced between these two time-points. These data suggest that the protection afforded by immunization with Vi-PS reduces slowly over time, but that even low titres of Vi-specific antibody are sufficient to reduce bacterial infection *in vivo*.

## Discussion

Except for toxins, capsular polysaccharide antigens are the only single-antigen vaccines used against bacterial infections and so understanding how capsular polysaccharides induce protection is important in the wider context of bacterial vaccinology. A major reason for the use of conjugate vaccines such as TCV is their ability to induce immune responses in infants. Nevertheless, on a broader level they are also useful tools to evaluate how immune responses differ to the same antigen dependent upon how it is encountered by the host – in the context of the current study as a conjugated or unconjugated antigen containing a limited number of carbohydrate epitopes. The differences in the outcomes from altering this presentation are profound, including a marked alteration in the persistence of the IgG antibody response and the frequency of plasma cells in niches such as the bone marrow. The recent licensure of the TCV and its success in challenge and field studies (21, 22, 24, 45), reports of hyporesponsiveness after Vi-PS (46-48) and the intention to examine the role of boosted immune responses on protection in children combine to make highlight the importance of studying the immunology of responses to different capsular polysaccharide vaccines. Such studies will help understand how to maximize the clinical benefit of TCV for both infants and adults across the life course.

Both unconjugated Vi-PS and TCV induced early IgM and IgG responses and classical extrafollicular responses; however, as expected, only mice immunized with TCVs induce germinal centre responses. Anti-Vi IgA may contribute to the protection conferred by Vi vaccines (49). In our studies, little IgA was detected and only consistently in the period shortly after immunization and so the role of IgA could not be studied here. Despite the differences in antibody responses to the conjugated and unconjugated vaccines, bacterial burdens in challenged, short-term immunized mice were similar, irrespective of vaccine type or number of doses given. In human challenge studies, similar vaccine efficacies were observed for both unconjugated Vi-PS and TCV vaccines in adult volunteers challenged one month after vaccination (50). Within that protective envelope, the immune mechanisms involved in protection may show some variance, reflecting the different types of responses induced to the T-dependent TCV and T-independent Vi-PS vaccines (49). Nevertheless, the fundamental finding from field and challenge human studies as well as our mouse studies is that Vi-based vaccines can induce significant protection that is dependent upon antibody and not whether this antibody is derived from germinal centre responses. Therefore, while the network of activities associated with the production of high-affinity antibodies to Vi antigen could potentially augment protection, this is not essential in adults. Therefore, germinal centre-derived antibody responses may provide other benefits here, such as promoting longer-lived responses. This overlap between the contribution of antibody type and persistence may complicate the identification of simple correlates of protection for Vi-based vaccines.

To test the concept that there was redundancy between different immunoglobulin isotypes for protection, we examined responses and protection after immunization with Vi-PS and TCV in mice lacking IgM or IgG. AID^-/-^ mice generated significant IgM responses to TCV. This reflects the size of the Vi polysaccharide used (average size 165 kDa (26)) being sufficiently large to induce T-independent responses (47), (28). The IgM induced in the first two weeks after immunization to both Vi-PS and TCV is similarly protective in the model systems used in the current study. This finding was consistent with IgM being sufficient and essential for protection and that IgG supplements this IgM-mediated protection (51, 52). Nevertheless, further studies showed that IgM is not essential to control infection by Vi-expressing bacteria. The simplest explanation as to why IgM and IgG are both able to provide independent protection is the surface-exposure of Vi antigen. The amounts of Vi on the surface can be sufficient to limit access of antibodies to other surface antigens like lipopolysaccharide (LPS) O-antigen (53-55). This indicates that antibody is able to access different Vi epitopes with relative ease, and there is little steric hindrance of Vi antigen epitopes by Vi antigen itself or by other antigens. This ease of access can help explain why there can be similar binding to epitopes by pentameric IgM or monomeric IgG. In other contexts, the interplay between antigens can restrict access of antibodies to bacterial antigens. For instance, LPS O-antigen can occlude access to protective epitopes in porin trimers by effectively shielding access and this can even influence the level of binding and protection afforded by different IgG isotypes (36, 56), independent of IgG effector functions. Thus, the surface exposure of Vi shields non-Vi antigens to antibodies but is itself accessible to Vi-specific antibodies of multiple isotypes, which may help explain why capsular polysaccharides make such excellent vaccine antigens.

The capacity of both IgM and IgG to moderate infection may also help explain why immunization with Vi-PS could still provide some protection months after immunization. Indeed, the maintenance of some protection after immunization with all vaccine types, despite differences in the levels of IgM and IgG and the isotypes induced, indicates that a key factor in protection was the persistence of antibody responses, rather than the nature of the antibody induced. In the context of using TCV in humans it can be hypothesised that TCV are superior to Vi-PS vaccines due to the longer lasting antibody responses induced and not the antibody isotypes induced. A single immunization with TCV offers excellent protection (57) but additional immunizations may enhance this protection through extending its longevity rather than its magnitude. In this regard it is noteworthy that in our studies, two immunizations of 2 µg Vi-TT induced higher IgG titres than one immunization with 4 µg Vi-TT. Moreover, consistent with what is presented here, an earlier study of anti-Vi responses and protection in children noted that boosting with Vi-rEPA results in high levels of protection that is not reduced by the halving of anti-Vi antibody levels over a two-year period (58).

The level of protection maintained in the Vi-PS-immunized mice at 7 months was surprising, with our challenge experiments indicating that immunization with 4 µg Vi-PS still provided some protection at this time. This suggests that further experiments are needed to identify vaccine doses that allow the study of long-term antibody responses induced to these vaccines and any waning of the protection they offer. Nevertheless, the 4 µg dose of Vi-PS used here is still substantially lower than that used in humans and mouse antibodies have a shorter half-life than their human equivalents (59). As such, the antibody response could reasonably have been expected to have fallen further from its peak than it had. For comparison, TyphimVi is administered at a dose of 25 µg (60), and trials have identified that the protection afforded by immunization with Vi-PS in adult humans wanes substantially by 2-3 years post-vaccination. Nevertheless, a difference in bacterial numbers in the liver was observed between 5 and 7 months after immunization with Vi-PS. This was not observed in mice immunized with Vi-TT and this probably is due to Vi-TT-boosted mice better maintaining their antibody responses. Collectively, these findings highlight the importance of understanding how persisting plasma responses to vaccines are maintained. This will require a deeper understanding of how germinal responses develop to vaccine antigens and of the niches in sites such as the bone marrow where long-lived plasma cells reside.

In conclusion, both Vi-PS and Vi-conjugate vaccines generate protective antibody responses that are not dependent upon only IgM or IgG. The results suggest that in contrast to Vi-PS, the ability to boost TCVs may provide superior protection by providing longer-lasting antibody responses.

## Acknowledgments

We gratefully acknowledge the staff of the Biomedical Services Unit at the University of Birmingham and the GSK Vaccines Institute for Global Health for their help and support. This research has received funding from the People Programme (Marie Curie Actions) of the European Union Seventh Framework Programme FP7/2007-2013/ under REA grant agreement 316940 awarded to the GSK Vaccines Institute for Global Health SRL (formerly Novartis Vaccines Institute for Global Health) and the University of Birmingham. SEJ received funding from the Midlands Integrative Biosciences Training Programme (MIBTP), a Doctoral Training Partnership of the Biotechnology and Biological Sciences Research Council (BBSRC) with additional funding from GVGH as part of a studentship programme. RRP received funding from the Wellcome Trust, and AA received funding from King Saud University.

## Author Contributions

SEJ and AFC wrote the manuscript. SEJ, MA, CAM, IRH, FM and AFC designed the studies. SEJ, MA, AA, RRP, EMJ, MPT, JP, WMC, AES, REL, CNC and MG performed the research and collected data. AJB and TH provided bacterial strains. LHJJ and KMT provided murine strains. EP, RDB and FM provided vaccine formulations. All authors critically revised and approved the final version of the manuscript.

## Competing Interest Statement

GSK Vaccines Institute for Global Health Srl is an affiliate of GlaxoSmithKline Biologicals SA. FM and RDB are employees of the GSK group of companies. SEJ and EP participate in a postgraduate studentship program at GSK. MA participated in a postgraduate studentship at GSK at the time of the study.

## Data Availability

All data are available in the main text or supplementary materials.

