## Supplemental figures for "Vi polysaccharide and conjugated vaccines afford similar early, IgM or IgG-independent control of infection but boosting with conjugated Vi vaccines sustains the efficacy of immune responses"

Supplementary figures

**A)**

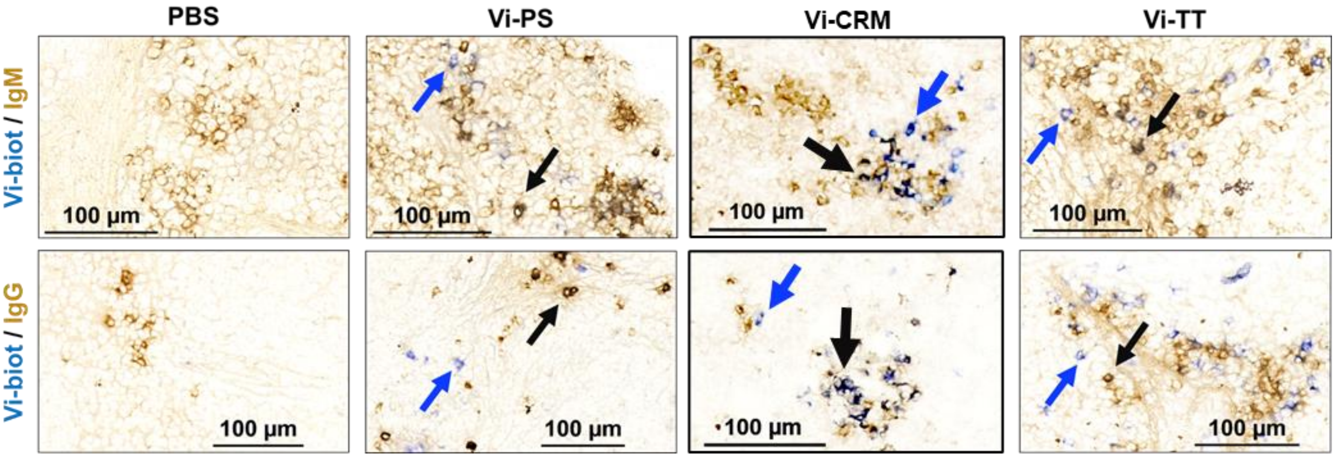


**B)**

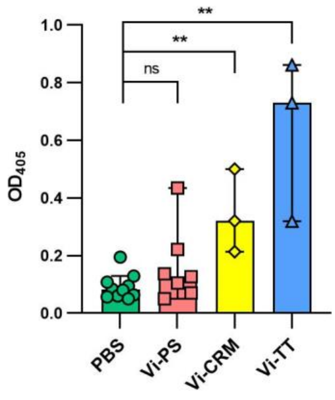


**Supplementary Figure 1. Immunohistochemical staining of splenic IgM and IgG positive Vi-specific cells day 7 after immunisation**. (A) Representative immunohistological images of spleen tissue from C57Bl/6 mice 7 days after *i.p* injection of 2 µg Vi-PS, Vi-$CRM_{197}$, Vi-TT or PBS. Spleens were stained for IgD, IgM or IgG (brown) and Vi (blue). Blue arrows = Vi positive, black arrows = double positive**. (B)** Anti-Vi IgA was detected by ELISA, reported as $OD_{405}$ values of serum at a 1:30 dilution. Bars represent median with 95% confidence intervals. Representative of 2 experiments with n = 3-6 mice/group. Bars represent median with 95% confidence intervals. ** = p≤0.01, and ns = non-significant by Mann-Whitney U test between individual groups (two-tailed).

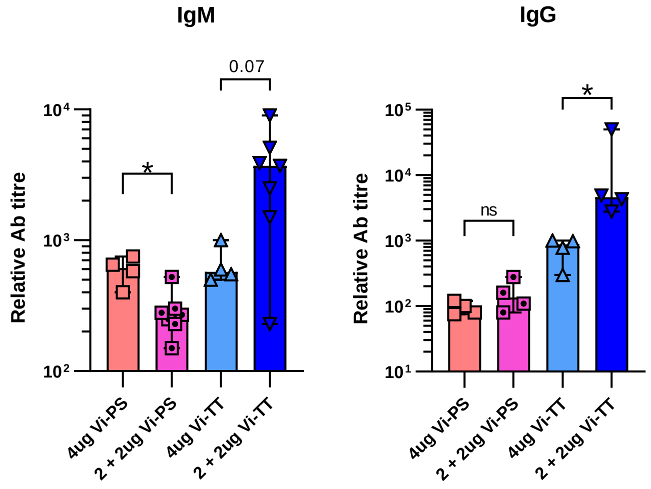


**Supplementary Figure 2. Vi-PS does not induce a booster response.** C57Bl/6 mice were immunized *i.p* with either 4 µg Vi-PS or Vi-TT on day 0, or 2 µg Vi-PS or Vi-TT on both day 0 and 35, then challenged with 1x10$^5$ CFU Vi+ *S.* Typhimurium TH177 from day 41-44. Sera were assessed by ELISA for anti-Vi IgM and IgG. Representative of 3 experiments with n = 2-4 mice/group. Bars represent median with 95% confidence intervals. * = p≤0.05, and ns = non-significant by Mann-Whitney U test between individual groups (two-tailed).

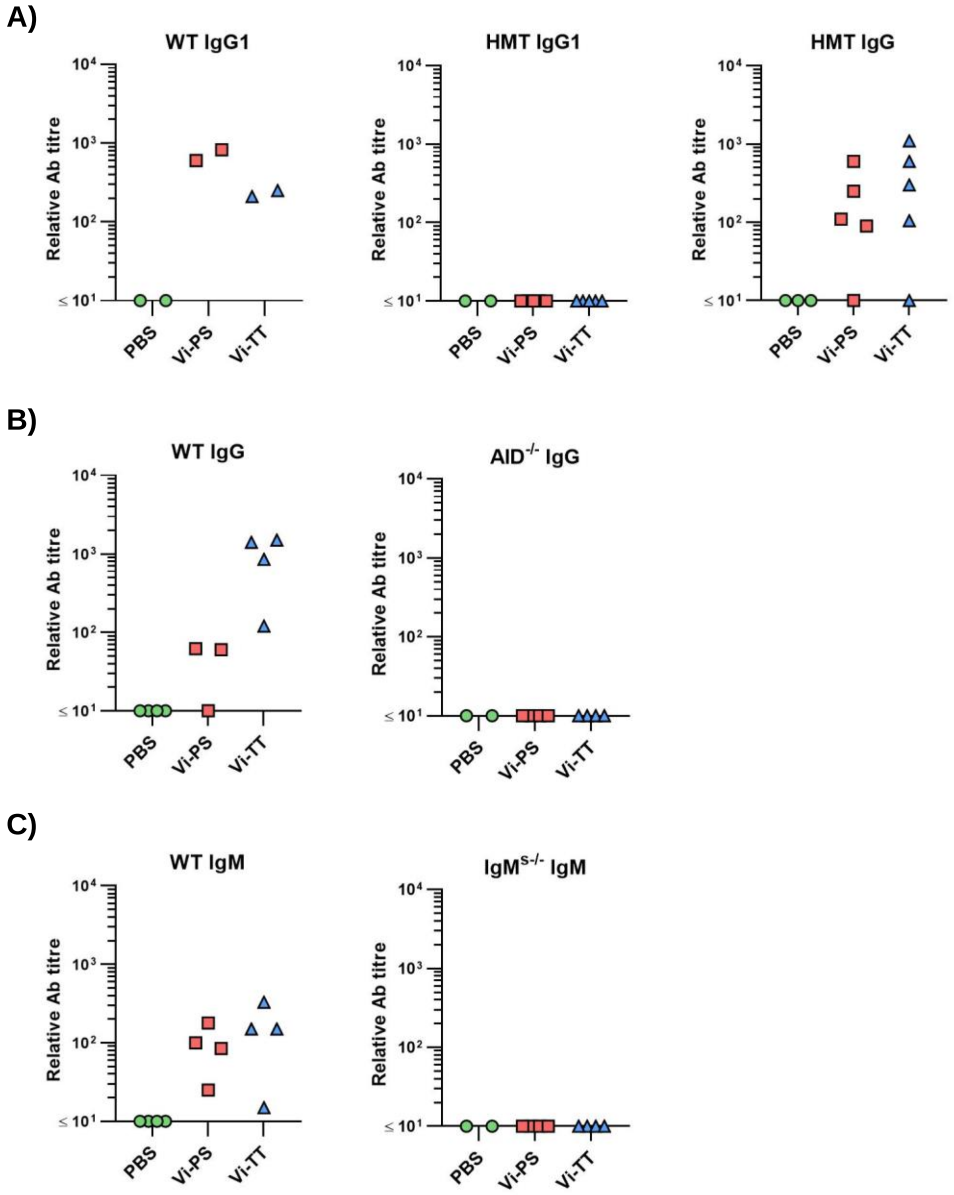


**Supplementary Figure 3. Serum antibody titres from Vi vaccine immunised antibody isotype knock out mice compared to WT mice.** C57Bl/6 (WT), IgG1$^{-/-}$ (HMT), AID$^{-/-}$ and IgM$^{s-/-}$ were immunised *i.p* with 2 μg Vi-PS, Vi-TT or PBS, then infected for 24 hours with 1x10$^{5}$ CFU Vi+ *S.* Typhimurium TH177 14 days later. Serum antibody was assessed by ELISA to show **(A)** Anti-Vi total IgG or IgG1 in WT compared to HMT mice, **(B)** Anti-Vi total IgG in WT or AID$^{-/-}$ mice or **(C)** Anti-Vi IgM in WT or IgM$^{-/-}$ mice. Each point represents the data from one mouse.

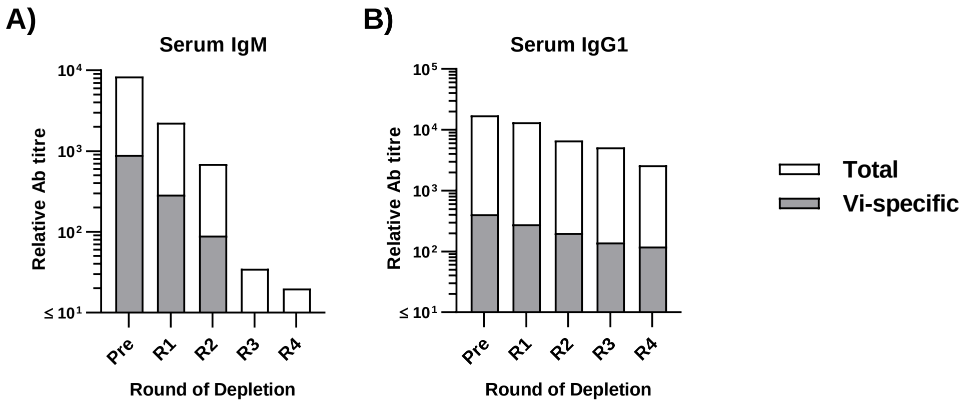


**Supplementary Figure 4. IgM depletion of Vi hyperimmune mouse serum.** Sera from 3 mice immunised at day 0, 14 and 35 with 2ug Vi-TT were collected day 42 post-primary immunisation. An equal volume of serum from each mouse was pooled. Pooled hyperimmune serum was then incubated with rat anti-mouse-IgM coated Sepharose beads to remove IgM four times. **(A)** After each round, serum was sampled and both total IgM (white bars), and Vi-specific IgM (grey bars) were assessed by ELISA. Compared to pre-depletion serum, 100% of Vi-specific IgM was lost. **(B)** Total and Vi-specific IgG1 was also assessed to confirm that depletion was specific to IgM. Compared to pre-depletion serum, 29% of Vi-specific IgG1 was lost through dilution.

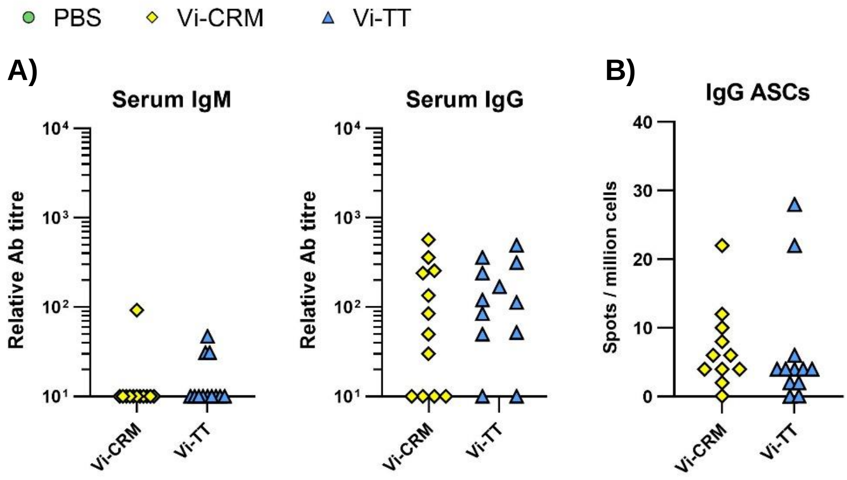


**Supplementary Figure 5. Antibody responses to a single dose of TCVs after 6 months.** C57Bl/6 mice were immunized *i.p* with 2 μg of Vi-$CRM_{197}$ or Vi-TT on day 0. Serum was collected day 185 (6 months post-primary immunization). Representative of 2 experiments with n = 5-6 mice/group. **(A)** Sera were assessed by ELISA for anti-Vi IgM and IgG. **(B)** Bone marrow anti-Vi IgG antibody secreting cells (ASCs) at the 6-month timepoint were enumerated by ELISPOT.

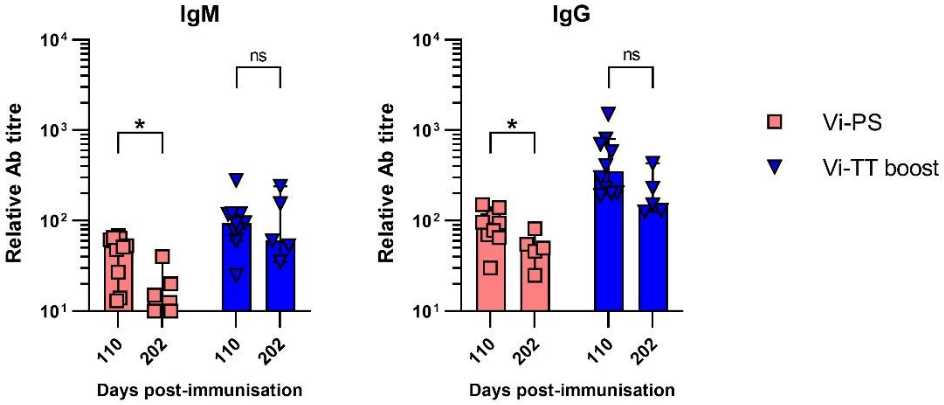


**Supplementary Figure 6. Change in antibody titres between day 110-202.** C57Bl/6 mice were immunised *i.p* with 4 μg Vi-PS on day 0 or 2 μg Vi-TT at day 0 and 35. Serum was collected at day 110 and 202 post-immunisation and anti-Vi IgM and IgG detected by ELISA. Representative of 2 experiments with n = 5-6 mice/group. Bars represent median with 95% confidence intervals. * = $p \leq 0.05$, and ns = non-significant by Mann-Whitney U test between individual groups (two-tailed).
